# A sensitized genetic screen to identify regulators of *C. elegans* germline stem cells

**DOI:** 10.1101/2021.10.03.462952

**Authors:** Sarah Robinson-Thiewes, Aaron M. Kershner, Heaji Shin, Kimberly A. Haupt, Peggy Kroll-Connor, Judith Kimble

## Abstract

Germline stem cells (GSCs) in *Caenorhabditis elegans* are maintained by GLP-1/Notch signaling from the niche and by a downstream RNA regulatory network. Loss of the GLP-1 receptor causes GSCs to precociously undergo meiotic differentiation, the “Glp” phenotype, due to a failure to self-renew. *lst-1* and *sygl-1* are functionally redundant direct targets of GLP-1 signaling whose gene products work with PUF RNA binding proteins to promote GSC self-renewal. Whereas single loss-of-function mutants are fertile, *lst-1 sygl-1* double mutants are sterile and Glp. We set out to identify genes that function redundantly with either *lst-1* or *sygl-1* to maintain GSCs. To this end, we conducted forward genetic screens for Glp mutants in genetic backgrounds lacking functional copies of either *lst-1* or *sygl-1*. The screens generated nine *glp-1* alleles, two *lst-1* alleles, and one allele of *pole-1*, which encodes the catalytic subunit of DNA polymerase ε. Three *glp-1* alleles reside in Ankyrin (ANK) repeats not previously mutated. *pole-1* single mutants have a low penetrance Glp that is enhanced by loss of either *lst-1* or *sygl-1*. Thus, the screen uncovered one locus that interacts genetically with both *lst-1* and *sygl-1* and generated useful mutations for further studies of GSC regulation.

## Introduction

Stem cells maintain a robust balance between self-renewal and differentiation to ensure tissue homeostasis despite physiological and environmental challenges. Failure to maintain that balance can lead to tissue dysfunction, disease, and death (Simons and Clevers 2011). Therefore, understanding the molecular circuitry governing stem cell regulation is critical. Yet biologically robust regulatory circuits are notoriously difficult to disentangle.

The *C. elegans* germline is a powerful system for the study of stem cell regulation (Hubbard and Schedl 2019; Gordon 2020). The adult hermaphrodite germline is contained in two U-shaped gonadal arms and produces oocytes; sperm are made during larval development and stored for later fertilization (Figure 1A, top). Germline stem cells (GSCs) are maintained at the distal end of each gonadal arm by a single-celled somatic niche, while GSC daughters differentiate as they move proximally away from the niche and ultimately undergo oogenesis (Figure 1A, middle)(Hubbard and Greenstein 2000).

**Figure 1:**
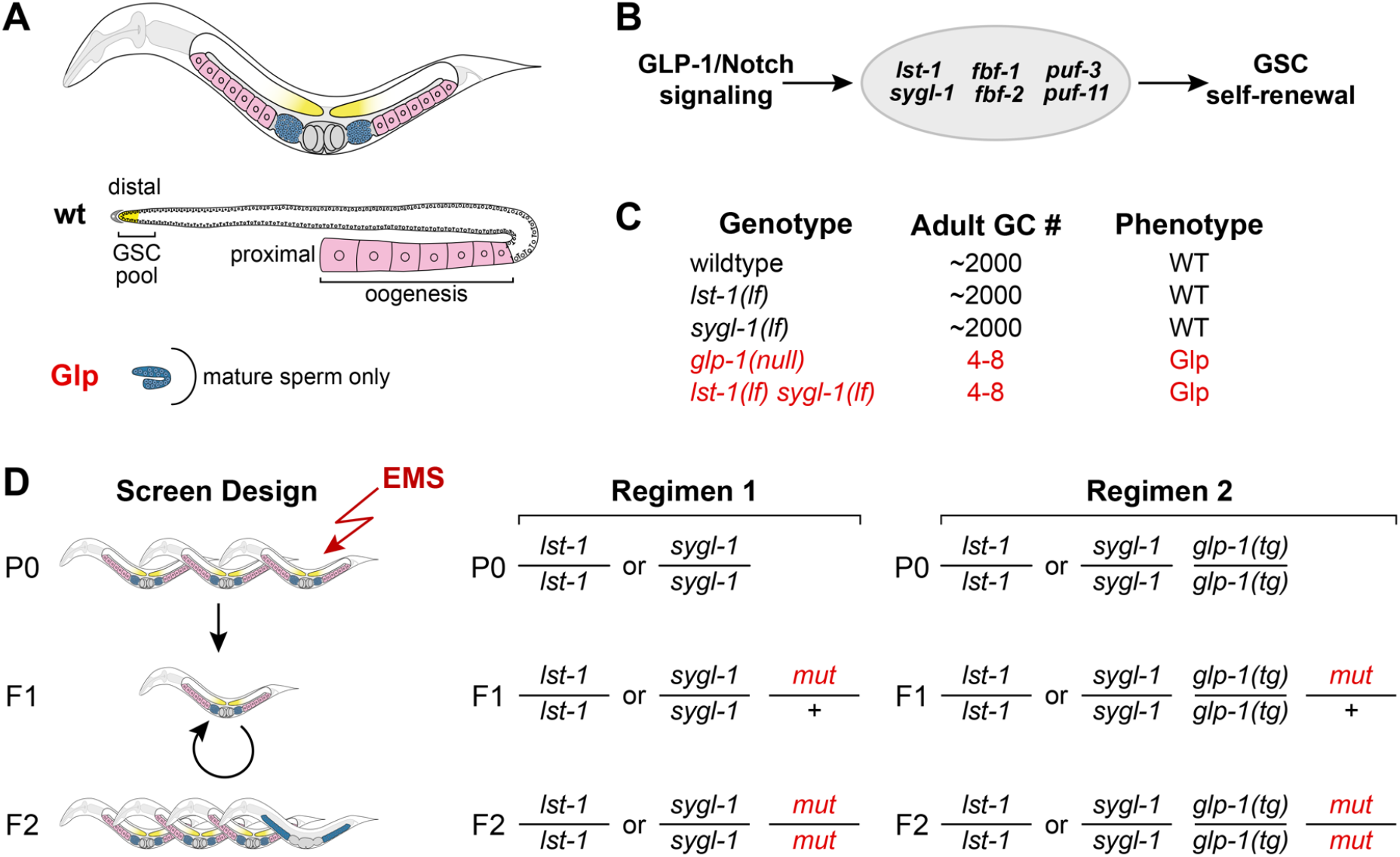
Genetic screens for synthetic Glp mutants. **A**. Top, adult hermaphrodite has two-U-shaped gonadal arms (GSCs, yellow; blue, sperm; pink, oocytes). Sperm made during larval development are stored in spermatheca. Middle, wildtype germline with a GSC pool (yellow) distally and oocytes (pink) proximally. Bottom, Glp adult germline with only a few mature sperm (blue). **B**. Molecular regulation of GSC self-renewal. GLP-1/Notch signaling activates transcription of *lst-1* and *sygl-1*, which are components of the PUF regulatory hub, along with *fbf-1, fbf-2, puf-3*, and *puf-11* (Haupt *et al*. 2019b). **C**. Adult germ cell (GC) numbers and phenotypes of specified genotypes. **D**. Strategies to identify genes that have a synthetic Glp phenotype with *lst-1* or *sygl-1*. Regimen 1 mutagenizes *lst-1(lf)* or *sygl-1(lf)* homozygotes and scores for Glp sterility in the F_2_. Regimen 2 mutagenizes *lst-1(lf)* or *sygl-1(lf)* homozygotes that also carry a wildtype *glp-1* transgene, *glp-1(tg)*, to avoid isolation of *glp-1* mutations.

GSC self-renewal depends on GLP-1/Notch signaling from the niche and on a downstream RNA regulatory network. In *glp-1* null mutants, GSCs fail to self-renew and instead differentiate precociously into sperm—the “Glp” phenotype (Austin and Kimble, 1987) (Figure 1A, bottom). Downstream of GLP-1/Notch, a “PUF hub” is required for self-renewal (Figure 1B). This regulatory hub comprises four genes encoding PUF RNA binding proteins as well as two direct GLP-1/Notch target genes, *lst-1* and *sygl*-1, that encode novel PUF interacting proteins (Crittenden *et al*. 2002; Kershner *et al*. 2014; Shin *et al*. 2017; Haupt *et al*. 2019a, 2019b; Qiu *et al*. 2019).

The PUF hub is characterized by pervasive genetic redundancy. For example, mutants lacking three PUF homologs are able to sustain some GSC self-renewing divisions, but animals lacking all four homologs phenocopy *glp-1* null mutants (Haupt *et al*. 2019b). Moreover, single mutants lacking *lst-1* or *sygl-1* are fertile and similar to the wildtype, while *lst-1 sygl-1* double mutants phenocopy *glp-1* null mutants (Figure 1C) (Kershner *et al*. 2014). The highly redundant nature of the PUF hub has hampered the identification of its component parts. Indeed, the LST-1 and SYGL-1 were not identified using standard forward genetic approaches, but instead were discovered using a candidate gene approach (Kershner *et al*. 2014), leaving open the possibility that additional components remain unidentified. For example, the LST-1 or SYGL-1 proteins might work with other unknown redundant factors. Here we describe the results of mutagenesis screens designed to identify regulators that function redundantly with *lst-1* or *sygl-1*.

## Methods

### Strain Maintenance

Unless noted otherwise, strains were maintained as previously described (Brenner 1974), at a temperature of 15°C. Balancers used to maintain recovered alleles were *hT2[qIs48]* (Siegfried and Kimble 2002) and *hIn1[unc-54(h1040)]* (Zetka and Rose 1992). Table 1 lists the strains used and their genotypes.

**Table 1:** Strains used in study.

| Strain | Genotype | Reference |
| --- | --- | --- |
| N2 | Wildtype | (Brenner 1974) |
| JK2877 | *unc-32(e189) glp-1(q175) III/ hT2[qIs48] (I; III)* | This work |
| JK4356 | *lst-1(ok814) I* | (Kershner *et al.* 2014) |
| JK4774 | *lst-1(ok814) sygl-1(tm5040) I/ hT2[qIs48] (I; III)* | (Kershner *et al.* 2014) |
| JK4899 | *sygl-1(tm5040) I* | (Kershner *et al.* 2014) |
| JK5135 | *sygl-1(tm5040) I; qSi44[Pglp-1::6XMYC::6xHIS::glp-1 3'end] II* | (Sorensen *et al.* 2020); This work |
| JK5203 | *lst-1(ok814) I; qSi44[Pglp-1::6MYC::6XHIS::glp-1 3'end] II* | (Sorensen *et al.* 2020); This work |
| JK5209 | *lst-1(q827) sygl-1(tm5040) I/ hT2[qIs48](I;III)* | This work |
| JK5277 | *lst-1(q826) I/ hT2[qIs48](I;III)* | (Shin *et al.* 2017) |
| JK5305 | *lst-1(q827) I/ hT2[qIs48](I;III)* | This work |
| JK5315 | *lst-1(q826) sygl-1(tm5040) I/ hT2[qIs48] I;III* | (Shin *et al.* 2017) |
| JK5606 | *lst-1(ok814) pole-1(q831) I /hln1 [unc-54(h1040)] I* | This work |
| JK5293 | *sygl-1(tm5040) pole-1(q831) I/hln1[unc-54(h1040)] I* | This work |
| JK5250 | *pole-1(q831) I/hln1[unc-54(h1040)] I* | This work |
| JK5268 | *pole(gk49) I/hln1[unc-54(1040)] I* | This work |
| JK5546 | *glp-1(q819) III/hT2[qIs48] (I; III)* | This work |
| JK5547 | *glp-1(q824) III/hT2[qIs48] (I; III)* | This work |
| JK5568 | *glp-1(q818) III/hT2[qIs48] (I; III)* | This work |
| JK5569 | *glp-1(q822) III/hT2[qIs48] (I; III)* | This work |
| JK5570 | *glp-1(q825) III/hT2[qIs48] (I; III)* | This work |
| JK5575 | *glp-1(q817) III /hT2[qIs48] (I; III)* | This work |
| JK5576 | *glp-1(q820) III/hT2[qIs48] (I; III)* | This work |
| JK5577 | *glp-1(q821) III/hT2[qIs48] (I; III)* | This work |
| JK5578 | *glp-1(q823) III/hT2[qIs48] (I; III)* | This work |

### Screen design and phenotype scoring

We screened for *lst-1* or *sygl-1* enhancers using a modified ethyl methanesulfonate (EMS) protocol (Brenner 1974). Fourth larval stage (L4) hermaphrodites were soaked in 25 mM EMS (Sigma: M0880) for 4 hours at room temperature, washed with M9, and placed on plates. F1 progeny were singled onto individual Petri dishes and allowed to self at 15°C. F2 adult progeny were scored for sterility by dissecting scope, and then L4 larvae were scored for a Glp phenotype using a Zeiss Axioskop compound scope equipped with DIC Nomarski optics, as described (Kershner *et al*. 2014). Each screen was done in two ways— first with single mutants *lst-1(ok814)* and *sygl-1(tm5040)* (Figure 1D, regimen 1) and then with each of the same mutants carrying a transgenic copy of wildtype *glp-1* (Sorensen *et al*. 2020) in addition to an endogenous copy of wildtype *glp-1* (Figure 1D, regimen 2).

### Allele identification

Following isolation of a Glp mutant, the starting *lst-1* or *sygl-1* allele was crossed away to test whether the Glp phenotype depended on loss of *lst-1* or *sygl-1*. Mutations were then mapped to a chromosome and tested for their ability to complement alleles of likely candidate genes. Mutants that were fertile as single mutants and mapped to chromosome I were tested for complementation with *lst-1(ok814) I*. Briefly, the double mutant (e.g. *mut-x sygl-1)* was balanced over the green balancer *hT2[qIs48]*, crossed to *lst-1(ok814) sygl-1(tm5040)/hT2[qIs48]* males, and non-green L4 male progeny (e.g. *mut-x sygl-1/ lst-1 sygl-1)* scored for Glp. Mutants that were sterile as single mutants and mapped to chromosome III were tested for complementation with *glp-1(q175) III*. Briefly, *unc-32 glp-1(q175)/ hT2[qIs48]* males were mated to each suspected *glp-1* allele and non-green male progeny scored for Glp. If an allele failed to complement either *lst-1* or *glp-1*, then Sanger sequencing was used to identify the molecular lesion. The *glp-1(q823)* allele was sequenced 2382 bp upstream of the 5’ UTR and 927 bp downstream of the 3’ UTR in addition to the exons and introns, but no lesion was found.

Whole genome sequencing was used to identify the likely lesion in *q831*, which was sterile as a single mutant and mapped to the right arm of chromosome I. Briefly, we picked ∼570 adult homozygotes, isolated DNA with Puregene Core Kit A (Qiagen ID: 158667) following manufacturer’s directions and submitted the DNA (∼100 ng) to the Wisconsin Biotechnology Core for sequencing using an Illumina MiSeq. The genome sequence was uploaded to a Galaxy server and analyzed by CloudMap, as previously described (Minevich *et al*. 2012). A premature stop codon occurred in one gene, *F33H2*.*5*, which resides on the right arm of chromosome I. *q831* failed to complement *F33H2*.*5 (gk49)* (Barstead *et al*. 2012), and the premature stop codon was confirmed by Sanger sequencing of DNA from *q831* homozygotes.

### Assay for temperature sensitivity of glp-1 alleles

Balanced strains carrying *glp-1* alleles were maintained at 15°C, 20°C, or 25°C for at least one generation before homozygous *glp-1* L4 progeny were scored for Glp.

### pole-1 phenotype assay

Homozygous *pole-1 (q831* or *gk49)* animals were distinguished from the balancer *hIn1[unc-54(h1040)]* by their kinked, uncoordinated movement. Homozygous mid-L4 hermaphrodites were raised at 20°C, anesthetized in levamisole, mounted on an agarose pad, and examined using a Zeiss Axioskop compound scope (Crittenden *et al*. 2017). Vulva formation—wildtype, multivulva, or vulvaless—was scored in addition to germline defects.

### Immunostaining

Strains were maintained at 20°C for immunostaining following published procedure (Crittenden *et al*. 2017). The SP56 polyclonal anti-sperm antibody (Ward *et al*. 1986), a gift from Susan Strome (UCSC, California), was diluted 1:200. The secondary antibody Alexa Fluor 555 donkey **α**-mouse (1:1000, Invitrogen #A31570) was added with DAPI (1 µg/mL) to mark DNA. Gonads were mounted in Vectashield (Vector Laboratories #H-1000), sealed with nail polish, and kept in the dark at 4°C until imaging.

### Microscopy

DAPI/SP56 stained gonads were imaged with a Zeiss Axioskop compound microscope equipped with a Hamamatsu ORCA-Flash4.0 cMos camera and a 63/1.4 NA Plan Apochromat oil immersion objective. Carl Zeiss filter sets 49 and 43HE were used for the visualization of DAPI and Alexa 555. Images were captured using Micromanager (Edelstein *et al*. 2010, 2014).

*lst-1 RNAi*

The *lst-1* RNAi clone from the Ahringer library (Fraser *et al*. 2000) was used. Briefly, *lst-1* RNAi or empty vector control (pL4440) containing HT115 bacteria were grown overnight at 37°C in 2xYT media containing 25 μg/μl carbenicillin and 50 μg/μl tetracycline. Cultures were concentrated, seeded onto Nematode Growth Medium (NGM) plates containing 1mM IPTG, then induced overnight. L4 hermaphrodites were fed, allowed to self, and progeny were scored for the Glp phenotype by DIC.

### GLP-1 protein conservation

Protein sequences for *C. elegans glp-1* orthologs from other *Caenorhabditis* species were acquired from Wormbase. Sequences of the ANK repeats were aligned using M-Coffee to examine amino acid conservation (http://tcoffee.crg.cat/apps/tcoffee/do:mcoffee) (Notredame *et al*. 2000).

## Results and Discussion

### Screens for Glp mutants in lst-1 and sygl-1 single mutant backgrounds

To identify new GSC regulators and perhaps new components of the PUF hub, we conducted genetic screens for mutations that cause a Glp phenotype in a *lst-1(lf)* or *sygl-1(lf)* single mutant background (Figure 1D). Our initial screens simply mutagenized *lst-1(lf)* and *sygl-1(lf)* single mutants and scored their F2 progeny for the Glp phenotype (Figure 1D, regimen 1). We screened 8749 haploid genomes after mutagenesis of *lst-1(lf)* and 5504 haploid genomes after mutagenesis of *sygl-1(lf)* (Table 2). This first set of screens recovered ten mutants. However, outcrossing revealed that Glp phenotypes did not depend on either *lst-1(lf)* or *sygl-1(lf);* therefore, these mutant alleles were not of genes functionally redundant with *lst-1* or *sygl-1*. Nine mutations, alleles *q817-q825*, caused a fully penetrant Glp phenotype and mapped to chromosome III (Table 3). Because the *glp-1* locus is large (∼7.4 kb) and located on chromosome III, these nine mutations were likely *glp-1* alleles. Indeed, all nine failed to complement *glp-1(null)* (Table 3). The 10^th^ allele *q831* caused a low penetrance Glp and was mapped to the right arm of chromosome I, at some distance from both *sygl-1* and *lst-1* loci. Therefore, this mutation must be a lesion in some other gene; its identity is described below.

**Table 2:** Summary of screens and alleles recovered.

| Parental Genotype<sup>1</sup> | Copies of *glp-1(+)*<sup>2</sup> | Number of haploid genomes screened | Glp mutants recovered<sup>3</sup> | Gene identities |
| --- | --- | --- | --- | --- |
| *lst-1(lf) I* | 2 | 8749 | 6 | 6 *glp-1* |
| *lst-1(lf) I; qSi44 II* | 4 | 7922 | 0 | n/a |
| *sygl-1(lf) I* | 2 | 5504 | 4 | 3 *glp-1*<br>1 *pole-1* |
| *sygl-1(lf) I; qSi44 II* | 4 | 3868 | 2 | 2 *lst-1* |

<sup>1</sup> Alleles were *lst-1(ok814)* and *sygl-1(tm5040)*.
<sup>2</sup> Animals without qSi44 have two endogenous copies of *glp-1(+)*. Animals with qSi44 have two endogenous and two transgenic copies of *glp-1(+)*.
<sup>3</sup> Mutants with Glp phenotype—small germline and sperm to distal end (Austin and Kimble 1987)

**Table 3:** Genetic characterization of sterile mutants from screens.

| Allele | LG<sup>1</sup> | Glp<sup>2</sup> | Failure to complement<sup>3</sup> |
| --- | --- | --- | --- |
| *q817* | III | +++ | *glp-1(q175)* |
| *q818* | III | +++ | *glp-1(q175)* |
| *q819* | III | +++ | *glp-1(q175)* |
| *q820* | III | +++ | *glp-1(q175)* |
| *q821* | III | +++ | *glp-1(q175)* |
| *q822* | III | +++ | *glp-1(q175)* |
| *q823* | III | +++ | *glp-1(q175)* |
| *q824* | III | +++ | *glp-1(q175)* |
| *q825* | III | +++ | *glp-1(q175)* |
| *q826* | I | — | *lst-1(ok814)* |
| *q827* | I | — | *lst-1(ok814)* |
| *q831* | I | + | *pole-1(gk49)* |

<sup>1</sup> LG, linkage group
<sup>2</sup> +++ , 100% penetrance; +, ~30% penetrance; —, not Glp as single mutants.
<sup>3</sup> Allele used in complementation test.

The initial screens were heavily biased for the recovery of *glp-1* alleles. To limit the isolation of more *glp-1* alleles, we introduced a transgenic copy of wildtype *glp-1* into the *lst-1(lf)* and *sygl-1(lf)* single mutants (Figure 1D; Table 2). The *glp-1* transgene, *qSi44* or *glp-1(tg)*, is a single copy insertion of wildtype *glp-1* on chromosome II that rescues a *glp-1* null mutant (Sorensen *et al*. 2020). Using the same EMS mutagenesis procedure as before, we screened 7922 *lst-1(lf); glp-1(tg)* haploid genomes and 3868 *sygl-1(lf); glp-1(tg)* haploid genomes. No Glp mutants were isolated from *lst-1(lf); glp-1(tg)* but two were recovered from *sygl-1(lf); glp-1(tg)* (Table 2). These mutations were subsequently determined to be alleles of *lst-1* (see below). Table 3 summarizes the genetic characterization of alleles recovered from the screen, and Table 4 summarizes their molecular lesions. Our failure to recover *sygl-1* alleles in the *lst-1(lf)* background shows that our screens were not performed to saturation. However, we note that the *sygl-1* locus is relatively small (621 bp coding region) and therefore likely a poor mutagenesis target.

**Table 4:** Molecular lesions in alleles recovered from the screen.

| Gene(allele) | Type of Mutation | Nucleotide Change | Codon change<sup>1</sup> | Amino acid change<sup>1</sup> |
| --- | --- | --- | --- | --- |
| *glp-1(q817)* | missense | C → T | CCG → UCG | P1111S |
| *glp-1(q818)* | nonsense | C → T | CAA → UAA | Q98Stop |
| *glp-1(q819)* | missense | C → T | CAU → UAU | H1000Y |
| *glp-1(q820)* | missense | T → A | AAU → AAA | N992K |
| *glp-1(q821)* | nonsense | C → T | CGA → UGA | R499Stop |
| *glp-1(q822)* | nonsense | T → G | UAU → UAG | Y176Stop |
| *glp-1(q823)*<sup>2</sup> | unknown | not found | n/a | n/a |
| *glp-1(q824)* | substitution | AC → CA in intron 4<sup>3</sup> | n/a | n/a |
| *glp-1(q825)* | splice site | G → A | n/a | n/a |
| *lst-1(q826)* | nonsense | C → T | CGA → UGA | R114Stop |
| *lst-1(q827)* | splice site | G → A | n/a | n/a |
| *pole-1(q831)* | nonsense | G → A | UGG → UGA | W1899Stop |

<sup>1</sup> n/a, not applicable
<sup>2</sup> see methods for more details
<sup>3</sup> 184 bp from 5' splice site

### Characterization of lst-1 alleles

The *lst-1* locus generates two RNA isoforms – one longer, called *lst-1L*, and one shorter, called *lst-1S* (Figure 2A; Table 4). Most *lst-1* alleles available prior to this work were isolated in deletion screens (Kershner *et al*. 2014) or engineered by CRISPR/Cas9 gene editing (Haupt *et al*. 2019a). In addition, one allele from these screens was previously reported, the nonsense mutant *lst-1(q826)*(Shin *et al*. 2017). Here we report a second allele obtained in the screen, *lst-1(q827)*, which alters the 5’ splice site in *lst-1L* intron 2 (Figure 2A; Table 4). As previously reported for *lst-1(q826), lst-1(q827)* was confirmed by complementation tests and Sanger sequencing. Both alleles are phenotypically similar to previously characterized *lst-1(lf)* mutants: as a single mutant, they appear wildtype and as *lst-1 sygl-1* double mutants they are Glp. These *lst-1* alleles will prove useful in future studies focused on *lst-1* function.

**Figure 2:**
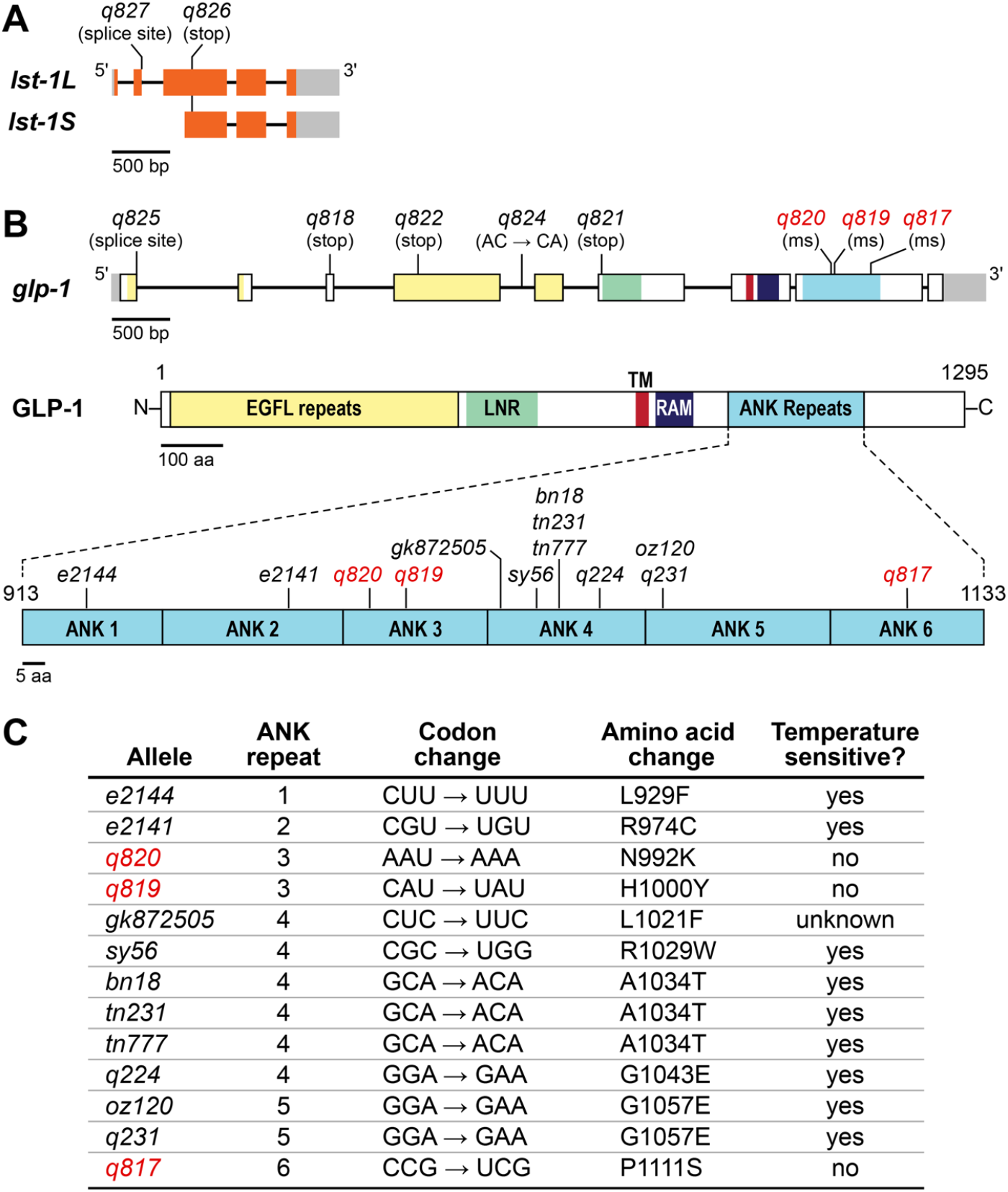
*lst-1* and *glp-1* alleles recovered from screens. **A-B.** Architecture of *lst-1* and *glp-1* loci. Boxes, exons with untranslated regions in gray; introns, lines connecting exons. **A**. The *lst-1* locus generates two RNA isoforms, *lst-1L* and *lst-1S*. Mutations isolated in screens shown above; see Table 4 for molecular changes. **B**. The *glp-1* locus generates one RNA isoform and one protein product. Regions within exons are colored according to protein domains: yellow, EGF-like (EGFL) repeats; green, *lin-12/Notch* Repeats (LNR); red, transmembrane domain (TM); dark blue, RAM domain; light blue, Ankyrin (ANK) repeats. Mutations in the ANK repeats that are shown below include those from this work (red) and those published previously (Austin and Kimble 1987; Kodoyianni *et al*. 1992; Berry *et al*. 1997; Dalfo *et al*. 2010; Thompson *et al*. 2013). Not shown are ANK repeat mutations isolated as intragenic suppressors of *glp-1(q231)* and *glp-1(q224)*(Lissemore *et al*. 1993). Ms, missense. **C**. Key features of *glp-1* mutations in ANK repeats. See Table 4 for molecular changes in other *glp-1* alleles and Table 5 for temperature sensitivity data.

### Characterization of glp-1 alleles

We identified the molecular lesions in the *glp-1* alleles with Sanger sequencing: *q818, q821*, and *q822* were nonsense mutants; *q817, q819*, and *q820* were missense mutants and *q825* altered a 5’ splice site (Figure 2B; Table 4). The *q824* allele had a 2 bp change (AC→CA) in intron 4 that did not affect the 5’ or 3’ splice sites or the branch point (Figure 2B). We failed to determine the lesion in one allele, *q823*, despite sequencing all exons and introns plus 2382 bp upstream of the transcription start site and 927 bp downstream of the 3’ UTR. Nonetheless, the remaining eight alleles were all previously unreported *glp-1* lesions.

The three *glp-1* missense alleles—*q817, q819*, and *q820* – all carry amino acid changes in the Ankyrin (ANK) repeats (Figure 2B and 2C). ANK repeats are conserved across eukaryotes with roles in protein interaction, cell signaling, and disease (Roehl *et al*. 1996; Mosavi *et al*. 2004). Many previously identified *glp-1* alleles also have changes in this region. Mutations in ANK repeats 1, 2, 4, and 5 all cause a temperature sensitive Glp phenotype (Kodoyianni *et al*. 1992; Berry *et al*. 1997; Nadarajan *et al*. 2009; Dalfo *et al*. 2010). Our three newly identified missense alleles occur in different repeats, ANK 3 (*q819* and *q820*) and ANK 6 (*q817*) and they are not temperature sensitive (Table 5). All three affect conserved residues (Figure 3). We conclude that the newly identified ANK missense mutations affect residues essential for GLP-1 function. These alleles should prove useful for investigating ANK repeats and their role in Notch signaling.

**Table 5:** *glp-1* alleles and temperature sensitivity.

| Allele | % Glp 25°C | % Glp 20°C | % Glp 15°C | n<sup>1</sup> |
| --- | --- | --- | --- | --- |
| N2 | 0 | 0 | 0 | 20 |
| *q175* | 100 | 100 | 100 | 20 |
| *q817* | 100 | 100 | 100 | 40<sup>2</sup> |
| *q818* | 100 | 100 | 100 | 20 |
| *q819* | 100 | 100 | 100 | 40 |
| *q820* | 100 | 100 | 100 | 40 |
| *q821* | 100 | 100 | 100 | 20 |
| *q822* | 100 | 100 | 100 | 20 |
| *q823* | 100 | 100 | 100 | 20 |
| *q824* | 100 | 100 | 100 | 20 |
| *q825* | 100 | 100 | 100 | 20 |

<sup>1</sup> n, number germlines scored.
<sup>2</sup>For *q817* at 15°C, 38 germlines scored.

**Figure 3:**
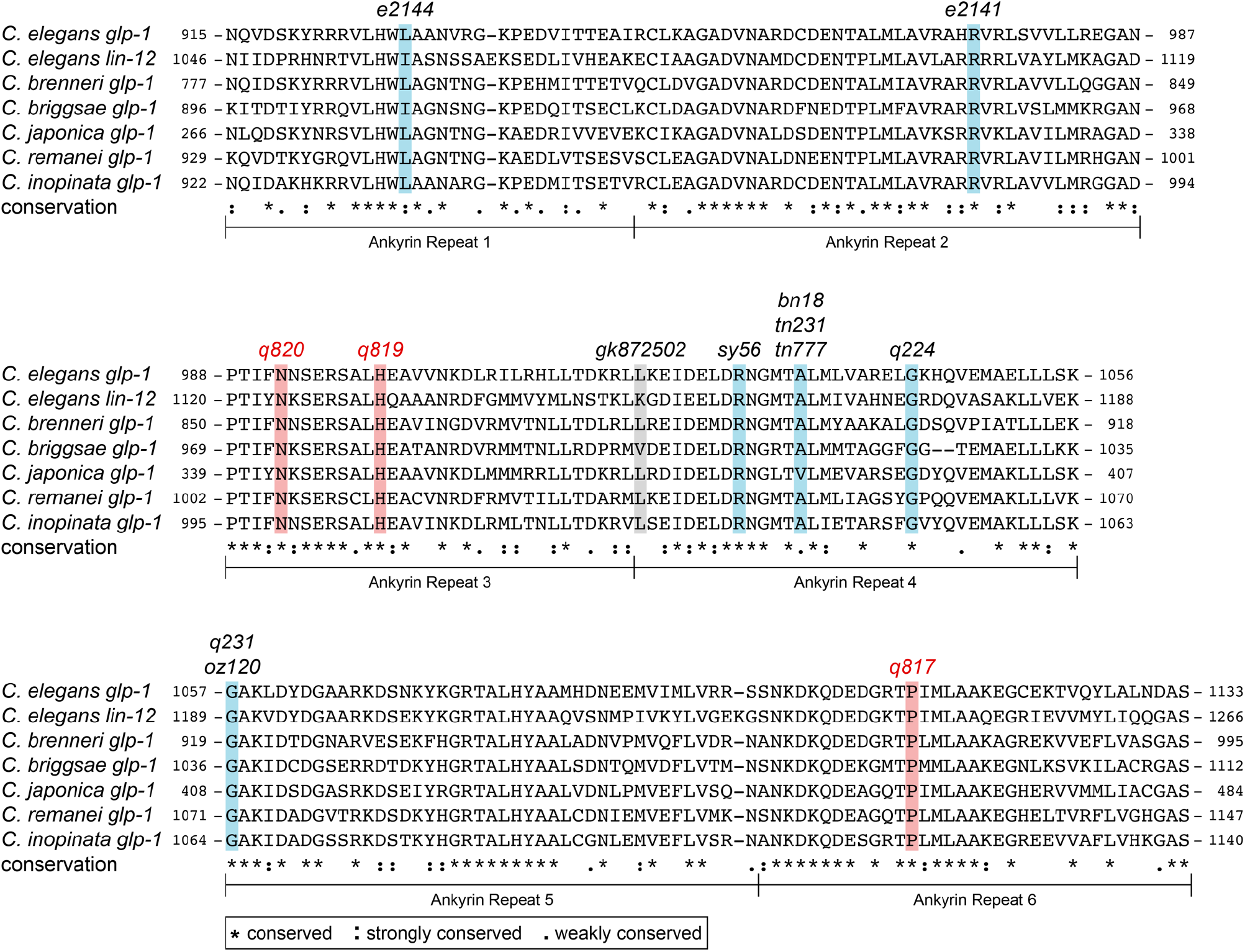
Amino acid alignment for ANK repeats *glp-1* orthologs and in the paralog *lin-12*. Alleles from Figure 2C are marked. Blue bar, mutation causes sterility at 25°C but not at 15°C; red bar, mutation causes sterility at 15°C, 20°C and 25°C. The residue affected in *gk872502* is marked by a gray bar, because it has not been tested for temperature sensitivity. ANK repeat location within each paralog is shown beside amino acids. See legend for conservation key.

### Characterization of pole-1(q831)

One mutant allele isolated in the *sygl-1(lf)* background, *q831*, mapped to the right arm chromosome I. Whole genome sequencing revealed a nonsense mutation R1899Stop in *F33H2*.*5* (Table 4), which encodes a *C. elegans* ortholog of the catalytic subunit of DNA polymerase ε (Figure 4A). We confirmed *q831* as an allele of *F33H2*.*5* by Sanger sequencing, and by its failure to complement *gk49*, a deletion allele in *F33H2*.*5* that had been generated by the *C. elegans* Knockout Consortium (Barstead *et al*. 2012). *F33H2*.*5* has been named *pole-1* for its DNA polymerase ε orthology.

**Figure 4:**
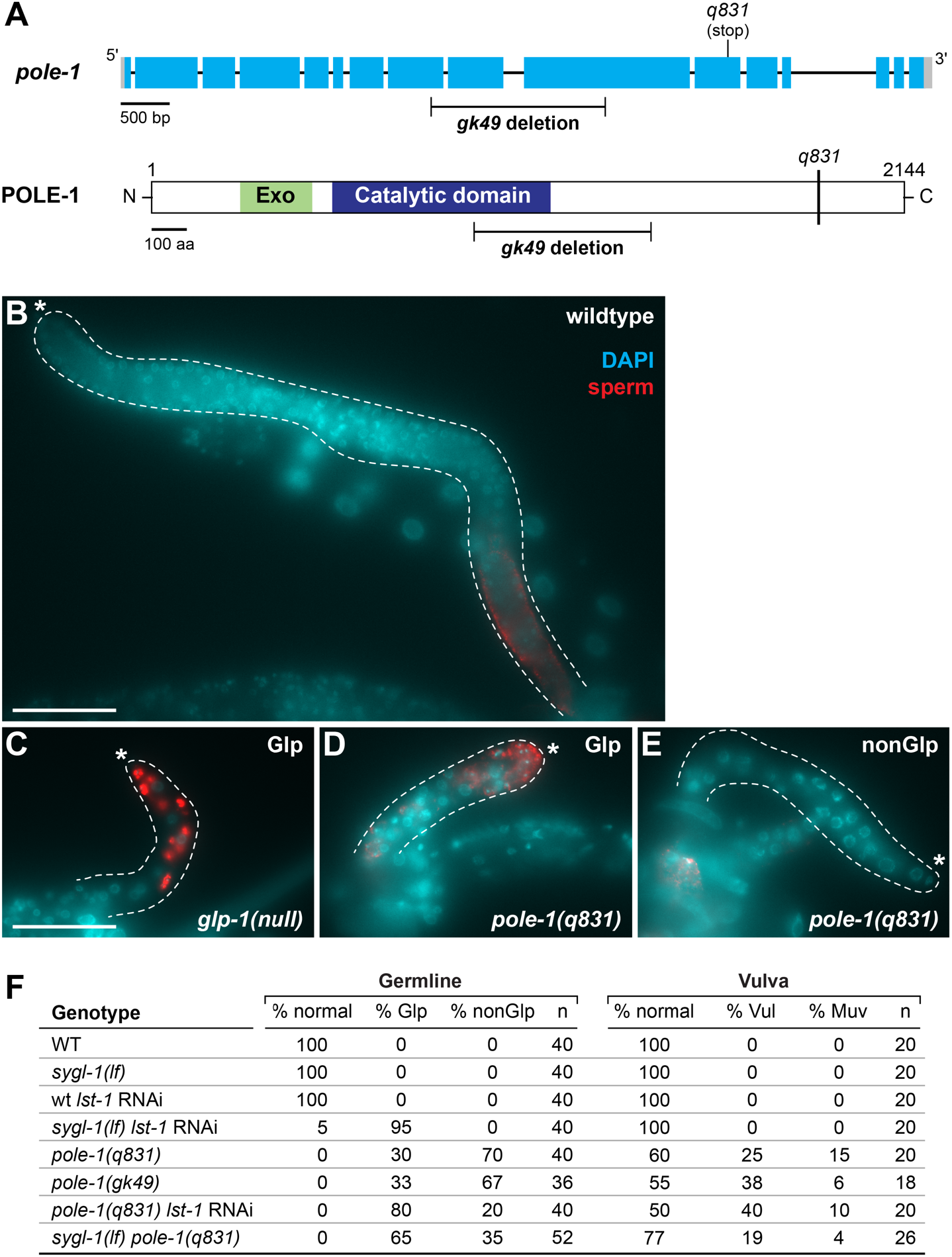
*pole-1* characterization. **A**. Diagrams of *pole-1* RNA and protein structures. Marked mutations include *gk49* (Barstead *et al*. 2012) and *q831* (this work). Conventions for gene structure as in figure 2. Protein domains: exonuclease (Exo) domain, green; DNA polymerase ε catalytic domain, dark blue (Pospiech and Syväoja 2003). **B-E**. Dissected mid-L4 gonads stained with SP56 antibodies for sperm (red) and with DAPI for DNA (blue) (see Methods). Dotted line outlines each gonad; asterisk marks the distal end. Scale, 50 µm. **B**. Wildtype. **C**. *glp-1* Glp germline. **D**. Glp *pole-1(q831)* germline. **E**. NonGlp *pole-1(q831)* germline. **F**. Low penetrance *pole-1* Glp phenotype is enhanced by loss of either *lst-1* and *sygl-1*. Germline” “normal” refers to an adult germline similar to wildtype in size and organization; “Glp” refers to a smaller than normal germline with sperm to distal end; “nonGlp” refers to a smaller than normal germline without sperm at the distal end. Vulva: “normal” refers a vulva similar to a wildtype morphology; “Vul” denotes Vulvaless; “Muv” denotes Multivulva. n, number of germlines or vulvas scored.

The *pole-1(q831)* mutation was isolated because *sygl-1(lf) pole-1(q831)* double mutants were Glp. During outcrossing, we found that *pole-1(q831)* single mutants were 100% sterile (Figure 4D-F). To ask if *pole-1* sterility was due to a Glp defect, we examined L4 larvae under DIC/Normaski and also stained dissected gonads with a sperm-specific antibody (SP56) (Ward *et al*. 1986) and DAPI (Figure 4B-F) (see Methods). Wildtype L4 gonads contain several hundred germ cells, with undifferentiated cells at the distal end and differentiated sperm at the proximal end (Figure 4B). *glp-1(null)* L4 gonads, by contrast, contain only a few germ cells, all of which have differentiated into SP56-positive sperm extending to the distal end (Figure 4C). Similar to *glp-1(null)* gonads, the *pole-1(q831)* gonads were physically smaller than wildtype; however only ∼30% had differentiated sperm extending to the distal end and thus were Glp (Figure 4D and F). The other ∼70% did not have sperm extending to the distal end and were designated nonGlp steriles (Figure 4E and F). We also observed a low penetrance Glp phenotype in the deletion strain *pole-1(gk49)*(Figure 4A, 4F). In addition to germline defects, *pole-1* mutants had a range of other defects, consistent with a broad role in development. For example, *pole-1* mutants had vulval defects (Figure 4F) and were uncoordinated.

We next asked if the *pole-1* Glp phenotype was enhanced by loss of either *lst-1* or *sygl-1*. Whereas *pole-1(q831)* single mutants were 30% Glp, *pole-1(q831) lst-1(RNAi)* animals were 80% Glp and *pole-1(q831) sygl-1(lf)* double mutants were 65% Glp (Figure 4F). Thus, loss of either *lst-1* or *sygl-1* enhanced the *pole-1* Glp defect. However, *pole-1* vulval defects were not similarly enhanced (Figure 4F). DNA polymerase ε *pole-1* had not been recognized as having an effect on GSC regulation though other components of the DNA replication machinery have been implicated in germ cell proliferation (Yoon *et al*. 2018). We conclude that *sygl-1* and *lst-1* are germline enhancers of *pole-1*.

## Conclusions and future directions

The goal of the mutant screens in *lst-1* and *sygl-1* mutant backgrounds was to identify new regulators of GSC self-renewal. In particular, we sought to test the idea that the LST-1 and SYGL-1 proteins might work with other factors that were similarly redundant. The screens identified nine alleles of *glp-1*, two alleles of *lst-1* and one allele of *pole-1—*the *C. elegans* ortholog of DNA polymerase ε. Although the screens were not saturated, identification of *pole-1* with a low penetrance Glp phenotype demonstrates that additional genes likely await discovery. Any additional screens in *lst-1* or *sygl-1* mutant backgrounds should focus on the modified design with transgenic *glp-1* to avoid isolation of more *glp-1* alleles. Alternatively, one might seek suppressors of *lst-1* or *sygl-1* tumors (Shin *et al*. 2017) or enhancers of the low penetrance *pole-1* Glp phenotype.

## Acknowledgements

We thank past and present members of the Kimble and Wickens labs for thoughtful discussions during the screens. We thank Erika Sorensen for sharing *glp-1(tg)* prior to publication, and Jadwiga Forster for technical support. The *gk49* allele was provided by the CGC, which is funded by NIH Office of Research Infrastructure Programs (P40 OD010440). SR-T was supported by the NSF Graduate Research Fellowship under Grant DGE-1256259 and NIH Predoctoral Training Grant in Genetics 5T32GM007133. JK was an Investigator of the Howard Hughes Medical Institute and is now supported by NIH R01 GM134119.

## Author Contributions

A.K, H.S, K.H and J.K designed screens and methods for mutant characterization; A.K, H.S, K.H., PK-C and J.K. performed screens; H.S. and K. H. characterized *lst-1* alleles; SR-T characterized *glp-1* alleles; A. K. and SR-T characterized *pole-1* alleles; SR-T, A.K, H.S, K.H, and J.K wrote the paper.

